# On the Etiology of Aesthetic Chills: a Behavioral Genetic Study

**DOI:** 10.1101/2021.07.08.451681

**Authors:** Giacomo Bignardi, Rebecca Chamberlain, Sofieke T Kevenaar, Zenab Tamimy, Dorret I Boomsma

**Affiliations:** Max Planck School of Cognition, Stephanstrasse 1a, Leipzig, Germany; Netherlands Twin Register, Department of Biological Psychology, Vrije Universiteit Amsterdam, The Netherlands; Department of Psychology, Goldsmiths University of London

**Author notes:** Address for correspondence: Giacomo Bignardi, corresponding author, Max Planck School of Cognition, Stephanstrasse 1a, Leipzig, Germany.

## Abstract

Aesthetic chills, broadly defined as a somatic marker of peak emotional-hedonic responses, are experienced by individuals across a variety of human cultures. Yet individuals vary widely in the propensity of feeling them. These individual differences have been studied in relation to demographics, personality, and neurobiological and physiological factors, but no study to date has explored the genetic etiological sources of variation. To partition genetic and environmental sources of variation in the propensity of feeling aesthetic chills, we fitted a biometrical genetic model to data from 14127 twins (from 8995 pairs), collected by the Netherlands Twin Register. Both genetic and unique environmental factors accounted for variance in aesthetic chills, with heritability estimated at .36 ([.33, .39] 95% CI). We found females more prone than males to report feeling aesthetic chills. However, a test for genotype x sex interaction did not show evidence that heritability differs between sexes. We thus show that the propensity of feeling aesthetic chills is not shaped by nurture alone, but it also reflects underlying genetic propensities.

## Introduction

Aesthetic chills ^1^ are embodied emotional-hedonic responses evoked by, among others, experiences with music ^2^, poetry ^3^, videos ^4^, beauty in nature or art ^5^, or even by eloquent speeches ^6^. They are frequently self-reported by individuals during peaks of hedonic ^7–9^ and emotional experiences ^2,10–14^, such as sadness and happiness ^12,15^, being moved ^14^, feeling touched ^10^, and the sensation of awe ^6^.

An extensive body of research has documented the physiological and neurobiological correlates of aesthetic chills, although sample sizes tend to be small. Chills usually occur with somatic manifestations, with participants reporting sidewise sensation of thrills in the upper dorsal part of the neck, or in the spine and back ^5^, shivers down the spine ^16^, tingling sensations in the arm ^17^, and more general diffuse bodily reactions ^10^. Bodily reactions that are associated with concurrent dynamic peripheral changes, mainly by increases in phasic skin conductance, and changes in heart rate, occur before, during and after chill onset ^2,8,9,13,18–21^. Besides being associated with self-reported and somatic manifestations, aesthetic chills usually correlate with activity in brain regions that play a role in the representation of visceral and somatic states ^22^, such as the bilateral insula (Ins) ^3,7,23^ and the anterior cingulate cortex ^7^, as well with several other brain regions that overlap with general reward mechanisms in the basal ganglia ^3,7,9,23^, and in the orbitofrontal cortex ^7^, plus other areas such as the supplementary motor area ^7,23^, the thalamus, and the cerebellum ^7,21,23^.

While most humans across cultures seem to have the capacity to experience aesthetic chills ^1^, individuals vary widely in the intensity and frequency with which they experience them ^2,6,24,25^. To date there is some evidence that demographic, personality and neurophysiological differences can account for some of this variation. For example, one study reports older individuals are more prone to report chills than younger ones ^26^, and others suggest females are more prone to report chills than males ^15,27^. It is worth noting however that results on the association between demographic factors are inconsistent across studies, with the vast majority reporting no significant effects of age ^20,28^ or sex ^2,20,29–31^.

Additional explanations for individual differences in the propensity of feeling chills come from studies on personality differences. Individuals who score higher on Openness to Experience (OE) tend to experience more chills, as measured both by self-report ^24,25,31–33^ and physiological measures ^20,24^. However, as for demographic correlates, it is also worth noting that the consensus on the relationship between personality factors and chills is far from being unanimous, with some suggesting higher correlations between emotional-aesthetic components of OE and the propensity of feeling chills ^31^, and others suggesting cognitive components of OE to play a bigger role ^24^. The picture is further complicated by few studies that suggest other aspects of personality play a role too ^4,13,20^, but, contrary to the association with OE, such studies have rarely been replicated.

Variation in the frequency of experiencing chills has also been accounted for by functional brain differences. For example, data obtained from resting-state functional Magnetic Resonance Imaging (rs-fMRI) on 1000 subjects indicates that individuals more prone to experiencing chills have enhanced connectivity between the network composed by the Ins and the cinglulate cortex (anterior and posterior cingo-opercular network) with ventral default network, by the latter network with the anterior visual and the posterior temporal networks, by the lateral and the dorsal default networks, and by a decrease in functional connectivity between the cerebellum and the somatomotor cortex^34^. Further, preliminary evidence obtained on small samples of individuals, suggests that the individual tendency of experiencing chills also correlates with structural brain differences, such as higher tract volume between the superior temporal gyrus and both the anterior insula and the medial prefrontal cortex ^19^, and resting physiological arousal, such as higher resting state skin conductance level ^28^.

The etiological sources of variation in aesthetic chills, i.e., how much of the observed variation can be explained by genetic and environmental factors, is yet unknown. Such lack of knowledge about the etiology is not unique to aesthetic chills alone but is shared among many studies on aesthetics. To our knowledge, there are only a few empirical investigations addressing the etiological sources of variation underlying individual differences in aesthetic experiences/appraisal ^35–40^. These studies applied the Classical Twin Model (CTM) to distinguish genetic from environmental influences. Two studies from the 1970’s (Barron ^35,36^) focused on individual differences in aesthetic sensitivity –defined as the extent to which one individual’s aesthetic judgment is in line with the opinion of experts-for paintings and drawings in a small sample of twins. The authors found contradicting results, with non-trivial heritability estimates ranging from 55% to 67% in the first study ^36^, and trivial estimates in the latter ^35^. The most recent study, from Butkovic et al. ^37^ found 40% of individual differences in flow proneness from music –a subjective, pleasurable, and fully absorbing experience— to be explained by genetic factors. Studies from Zietsch et al. ^40^, Germine et al. ^38^, and Sutherland et al. ^39^ focused on aesthetic preferences for faces. Heritability estimates were 33% for specific preferences for dimorphic male traits ^40^, and 22% to 30% for more general individual preferences for faces respectively.

Here we aim to investigate whether genetic effects can account for individual differences in the propensity of feeling aesthetic chills. To partition genetic and environmental sources of variation, we fitted a biometric genetic model to twin data, exploring a genotype by sex interaction by testing for both quantitative and qualitative sex differences. This allowed to test for differences in the importance of genetic influences on aesthetic chills and to test whether the same genes are expressed in men and women. We analyzed Item 43 of OE, “Sometimes when I am reading poetry or looking at a work of art, I feel a chill or wave of excitement” ^41^ as a proxy for the propensity of feeling chills. This item was selected, because explicitly asking individuals if they feel chills is a good indicator of actual experienced chills measured in experimental settings ^11,23,24^. Item 43 seems capable of tapping into individual differences which are highly shared among cultures ^1^ and to capture chills measured both by self-report and physiological changes ^24^, as well as neurophysiological differences between individuals ^34^.

## Method

### 2.1 Participants

The data were obtained from the Netherlands Twin Register (NTR), a longitudinal cohort established in 1987 by the department of Biological Psychology at the Vrije Universiteit Amsterdam. Data on the self-reported propensity of feeling aesthetic chills were collected by mailed surveys (see ^42^ for details), collected in 2004 (NTR survey 7 ^43^) on 6760, in 2009 (NTR survey 8 ^44^) on 10176 and in 2013 on 9419 twins (survey 10 ^45^). After excluding 320 pairs, for which no data for the item 43 were available across surveys, and data from 5 twin pairs due to missing information on age for both twins, we analyzed data on 14127 twins (9466 females), ranging from 14 to 97 years old, with mean age = 30 (SD = 13). Our procedure for data selection when multiple surveys had been completed, is detailed below. Table 1 shows the numbers of twins and the number of complete pairs (i.e., pairs in which both twins completed each survey) or incomplete pairs (i.e., pairs for which only one of the two twins completed the survey). Zygosity in same-sex pairs was based on genotyping for part of the sample and on survey information for others; ^42^ Test-reliability correlations for the twins who have completed more than one survey (N= 6923) were calculated for males and females separately. To avoid confounding effects due to familiar resemblances, data from one randomly selected twin per family were analyzed.

**Table 1.** Sample (N) of monozygotic (MZ) and Dizygotic (DZ) twin pairs per survey.

| Survey | N pairs | MZ<br>male | MZ<br>female | DZ<br>male | DZ<br>female | DZ<br>opposite sex |
| --- | --- | --- | --- | --- | --- | --- |
| Survey 7 | 6195 | 931 | 2463 | 478 | 1137 | 1186 |
| Survey 8 | 9100 | 1239 | 3242 | 747 | 1684 | 2188 |
| Survey 10 | 8302 | 1161 | 2935 | 663 | 1505 | 2038 |
| **Combined** | **8995** | **1279** | **2809** | **813** | **1645** | **2449** |
|  | **(5132)** | **(780)** | **(2010)** | **(392)** | **(914)** | **(1036)** |

Note. Combined sample is shown in bold. Number of complete pairs is shown between parentheses.

### 2.2 Materials

Self-report of chills was obtained from the short version of the NEO-personality Five Factor Inventory (NEO-FFI, ^41^). The NEO-FFI consists of 60 items rated on a five-point scale (1–5, totally disagree, disagree, neutral, agree and totally agree). The Openness to Experience (OE) scale contains an item (43) “Sometimes when I am reading poetry or looking at a work of art, I feel a chill or wave of excitement” which was selected as a proxy for the propensity of experiencing aesthetic chills.

### 2.3 Procedures

For 49% of participants, we had more than one survey. To maximize sample size, data from survey 7, 8 and 10 were merged into one data file by randomized selection of twin pairs per survey. The selection of pairs followed a number of criteria: 1) we prioritized complete answers from twin pairs, i.e., twin pairs were selected when both twins reported scores for the Item 43 on one of the surveys; 2) if one of the two twin’s response was missing for all three surveys, we randomly selected a survey with the response for the other twin; 3) if twins took part in different surveys, we randomly selected data for the pair from one of the complete surveys. Of the reported combined sample size, 5132 were twin pairs who both completed the survey, of which 2790 were MZ, 1306 were same-sex DZ (DZss), and 1036 were opposite-sex DZ (DZos) twin pairs.

### 2.4 Genetic analysis

We analyzed the data using the CTM to estimate the proportion of variance explained by genetic and environmental factors. Within the CTM, the observed phenotypic variance (P) can be decomposed into additive genetic (A), dominance genetic (D), common environmental (C), or unique environmental (E) components ^46^. The A component captures all additive effects of alleles across genetic loci, while the D component captures non-additive and interactive effects of alleles at contributing genetic loci. For the environmental components, the C captures all environmental factors that are shared between twins. When twins are raised together, home-environment effects can be captured by C. The E component captures all factors that are non-shared between twins nor explained by genetic factors. Thus, the E component captures all the unexplained unsystematic variance, comprising measurement errors.

The decomposition of the variance is possible because different genetic association exist for MZ and DZ twins. Since MZ twins derive from the same fertilized egg their genetic material is ∼100% shared (but see ^47,48^). MZ twins thus share 100% of both additive as well as dominance genetic effects. DZ twins, on the other hand, derive from two fertilized eggs and share only 50% on average of the additive genetic effects, and only 25% of the dominance genetic effects (see ^49^). Thus, within the CTM, the correlation between the A component within MZ twin pairs is equal to 1, while within DZ twins is equal to 0.5. Similarly, the correlation between the D component within MZ twin pairs is also 1, while for DZ pairs is .25. Further, since the C component captures all shared environmental effects and the E component captures all non-shared effects, the correlation for both MZ and DZ twins are set to 1 for the C component and 0 for the E component. As a consequence of these premises, when MZ twins resemble each other more than DZ twins on a given trait the heritability of such trait is considered to be non-trivial. Here, it is important to note that a model with four components cannot be statistically identified within the CTM, given that there is only enough information to estimate three components. Therefore only A,C and E or A, D, and E components can be estimated simultaneously ^49^. A rule of thumb is to assess the twin correlations and fit an ACE model accordingly when the MZ correlation is not larger than twice the DZ correlation. In comparison, an ADE model is fitted when the MZ correlation is larger than double the DZ correlation. Components within a model (e.g., ACE) are then dropped (e.g., dropping C) to assess whether they contribute to phenotypic variation. If the model with fewer components (e.g., AE) fits the data as equally well as the model with more components, then the dropped component is evaluated to be not explaining any significant phenotypic variance.

To investigate sex differences in the etiology of the trait, we look into the difference in correlations between female, male, and opposite-sex twins. When the genotype affects phenotypic variation to the same extent in men and women (no quantitative sex differences), we expect MZ male-male correlations to be equal to MZ female-female correlations and DZ male-male, and female-female correlations also to be similar. When there is no difference in the expression of the genes in men and women (no qualitative sex differences), we expect that the correlation in same-sex DZ twins is equal to that of opposite-sex DZ twins. Under the assumption of no sex effects, the amount of phenotypic variance is expected to be accounted for by a similar amount of A, D, C, and E, and similar genes are expected to influence phenotypic variation in both sexes ^50^.

The models were specified in OpenMx version 2.17.2 ^51–53^, in R-Studio, R version 3.6.2. Significance of the covariate (age), birth order effect and mean differences across zygosity were tested by a series of models nested inside the saturated model. The goodness of fit of the model was evaluated by 1) likelihood ratio, that is by the difference in minus twice the value of the log-likelihood (−2LL) between the two models, which has a *χ*^2^ distribution, with the degrees of freedom equal to the difference in the number of parameters, and by 2) the Akaike’s Information Criterion (AIC), by keeping the model with the lowest AIC as the best fitting model.

Genotype x sex interaction was tested for both quantitative sex differences (i.e., is the amount of variance in the propensity of feeling aesthetic chills accounted for by the same genetic effects across sexes?) and qualitative sex differences (i.e., are genes influencing variation in the propensity of feeling aesthetic chills the same in females and males?) in etiology. First, similarly to Vink et al. ^50^, the significance of differences in means, variances, and covariances across sexes was tested by a series of models nested inside the saturated model. A sex-limitation model was evaluated to examine whether quantitative sources of etiological variation statistically differed between the sexes by constraining variance components to be equal for men and women. The goodness of the fit and the significance of the sub-models with variance components constrained to be equal across sexes were compared to the full model. Qualitative sex differences were tested by allowing genetic correlations (r_g_) between DZos to be freely estimated. The goodness of the fit and the significance were obtained by comparing the more parsimonious model in which r_g_ within DZos was constrained to .5. Mean, standard deviations, 95% CI, and within twin pair correlations were estimated in a saturated model. A variance decomposition model was compared to the saturated model. Subsequently, we test nested models, which were obtained by constraining one of the genetic or environmental variance components to zero. Heritability estimates were obtained as the proportion of genetic variance over the phenotypic variance.

## Results

### 3.1 Descriptive statistics

Test-retest reliability was obtained for data from surveys 7 and 8 (5 years apart), surveys 8 and 10 (4 years apart), and surveys 7 and 10 (9 years apart). Single-item reliability estimates range from *r*(1488) = .58 ([.54,.61] 95% CI) and *r*(578) = .61 ([.56,.66] 95% CI between survey 7 and survey 8 for female and male respectively, *r*(2118)= .58 ([.55,.60] 95% CI) and *r*(830) = .52 ([.47,.57] 95% CI) between survey 8 and survey 10 for female and male respectively, and *r*(1078) = .58 ([.52,.60] 95% CI) and *r*(410) = .51 ([.44,.58] 95% CI) between survey 7 and survey 10 for female and male respectively (all *p* < .001, after Bonferroni correction).

The distribution of the item 43 scores is given in Figure 1 for first and second born twins, separately by sex. Individuals scale point frequency on the 5 Likert-scale ranged from 21% (“strongly disagree”) to 4% (“strongly agree”), with the majority of individuals (75%) distributed within the three central scale points. The finding that 21% of the firstborn twins do not report aesthetic chills in the combined survey is in line with some previous random-population sampling studies on chills ^2,5,16,17,30^.

**Figure 1.**
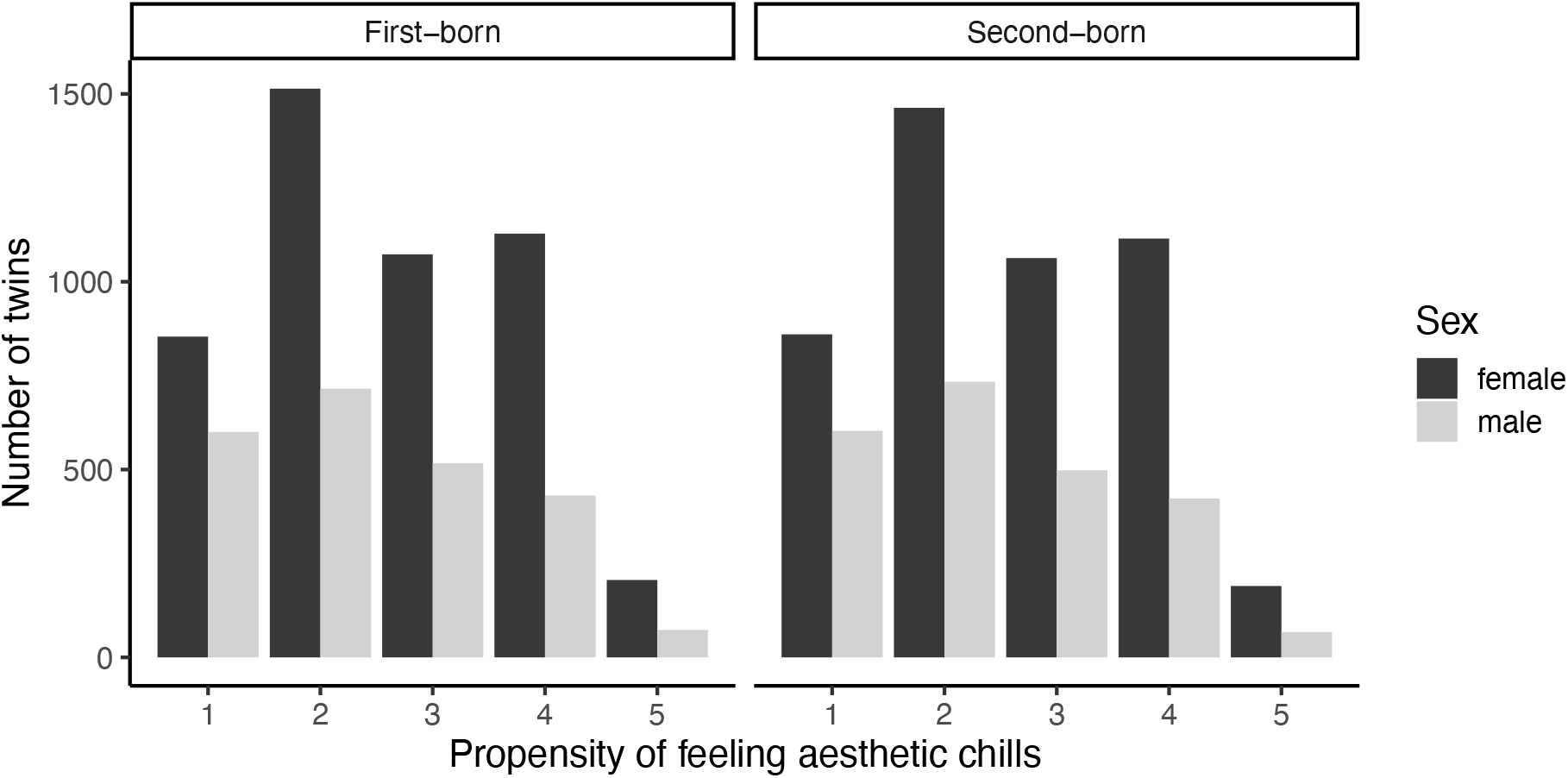
Distribution of item 43. “Sometimes when I am reading poetry or looking at a work of art, I feel a chill or wave of excitement”. The left panel shows the distribution for the first-born twin of the pair from female and male respectively. The right panel shows the distribution for the second-born twin of the pair from female and male respectively.

### 3.2 Biometric modelling

Twin correlations from the saturated model, were *r* = .39 ([.33,.44] 95% CI) and *r* = .35 ([.32,.39] 95% CI) for MZ male and MZ female respectively, *r* = .07 ([.00,.16] 95% CI) and *r* = .21 ([.14,.27] 95% CI) for DZ male and DZ female respectively, and *r* = .14 ([.08,.19] 95% CI) for DZos (also see Figure 2a). These correlations and CI suggested that an AE model to be most appropriate for describing these data. Table 2 shows the goodness of the fit comparison with the full saturated model. One the one hand, removing age as a covariate resulted in a deterioration of the model fit (−2LL = *χ*2(1) = *255*.*07, p* < .*001*). On the other, removing birth order and subsequently zygosity mean and variance differences did not deteriorate the overall model fit (all *p ≥* .*80*), indicating that mean and variance were not different across the first and the second-born and across zygosity. As expected, constraining mean scores to be equal across sexes resulted in a deterioration of the fit (−2LL = *χ*2(17) = *113*.*68, p* <.*001*), However, constraining variance to be equal across sexes, as well as constraining covariance to be equal across DZss and DZos, did not deteriorate the overall fit of the model (−2LL = *χ*2(17) = 6.64, *p =* .*99 and* -2LL = *χ*2(20) = 13.93, *p =* .*83*, respectively)

**Figure 2.**
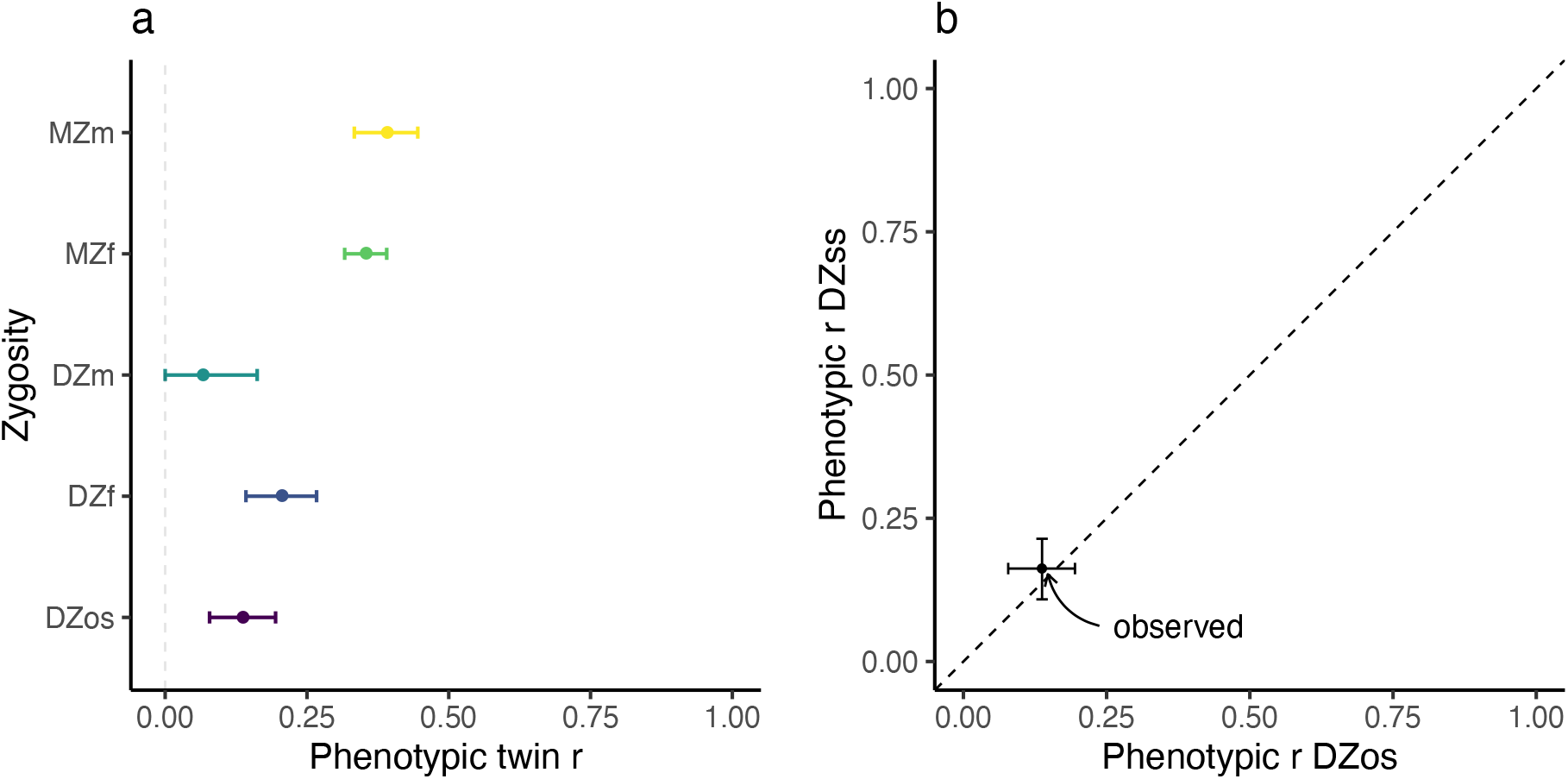
Phenotypic twin correlations. **a** Correlations (r) within twin pairs, error bars represent 95% CI. **b** adapted from Vink et al. ^50^. The dashed line represents the expected slope for the relationship between DZss and DZos r when genotype x sex interaction effects on phenotypic variation are not present. The dot represents the observed DZos pair correlation versus the DZss pair correlation, extracted from the Sex:DZss same covariance model. The horizontal and vertical error bars represent the 95% CI for the DZos and the DZss 95% CI. MZ = Monozygotic; DZ = Dizygotic; m = male; f = female; ss = same-sex; os = opposite-sex.

**Table 2.** Saturated model: Model-fitting results from five groups (MZ male, MZ female, DZ male, DZ female, DZos) model

| Model | -2LL | df | $\chi^2$ | $\Delta df$ | p | AIC |
| --- | --- | --- | --- | --- | --- | --- |
| Saturated Model | 43163.60 | 14101 | NA | — | — | 14961.60 |
| Covariate |  |  |  |  |  |  |
| *No age* | *43418.67* | *14102* | *255.07* | *1* | *<.001* | *15214.67* |
| Birth order |  |  |  |  |  |  |
| Same mean | 43165.21 | 14105 | 1.61 | 4 | .80 | 14955.21 |
| Same mean & variance | 43167.16 | 14109 | 3.55 | 8 | .89 | 14949.16 |
| Zygoty |  |  |  |  |  |  |
| Same mean | 43169.35 | 14113 | 5.75 | 12 | .92 | 14943.35 |
| Same mean & variance | 43170.22 | 14117 | 6.62 | 16 | .98 | 14936.22 |
| Sex |  |  |  |  |  |  |
| *Same mean* | *43277.28* | *14118* | *113.68* | *17* | *<.001* | *15041.28* |
| Same variance | 43170.24 | 14118 | 6.64 | 17 | .99 | 14934.24 |
| MZ same covariance | 43171.43 | 14119 | 7.823 | 18 | .98 | 14933.43 |
| DZss same covariance | 43177.14 | 14120 | 13.54 | 19 | .81 | 14937.14 |
| **DZ same covariance** | **43177.53** | **14121** | **13.93** | **20** | **.83** | **14935.53** |

Note. In bold best-fitting model. In Italics models that showed deterioration of the fit. Models are reclusively nested starting from the most parsimonious model. For example, the “‘Birth order: same mean & variance model’ is nested from the most parsimonious ‘Birth order: Same mean’, while ‘Birth order: Same mean’ is not nested in the ‘Covariate: *no age*’ model, since removing the covariate results in a deterioration of the overall fit. MZ = Monozygotic; DZ = Dizygotic; ss = same-sex; os = opposite-sex.

Phenotypic correlations for DZss and DZos, extracted from the model in which covariance across sexes were constrained to be equal (Sex:DZss same covariance model), were *r* = .16 ([.11,.21] 95% CI) and *r* = .14 ([.08,.19] 95% CI) respectively. The pattern of correlations between DZss and DZos indicates an absence of evidence of genotype x sex interaction effects (see Figure 2b). Table 3 shows the results for the sex limitation models. The full AE model was fitted to data from males and females, with separate estimates for means and variance components. As expected, the mean scale point for item 43 was found to differ across sexes (−2LL = *χ*2(1) = 106.32, *p* = <.001). However, constraining DZos r_g_ to be equal to .5 did not deteriorate the model fit (−2LL = *χ*2(1) = 2.16, *p =* .14). This indicated etiological sources of variation to not qualitatively differ across sexes. Moreover, constraining variance components across sexes to be equal did not deteriorate the model fit (−2LL = *χ*2(3) = 2.19, *p =* .53). This indicated etiological sources of variation to not quantitatively differ across sexes either.

**Table 3.**
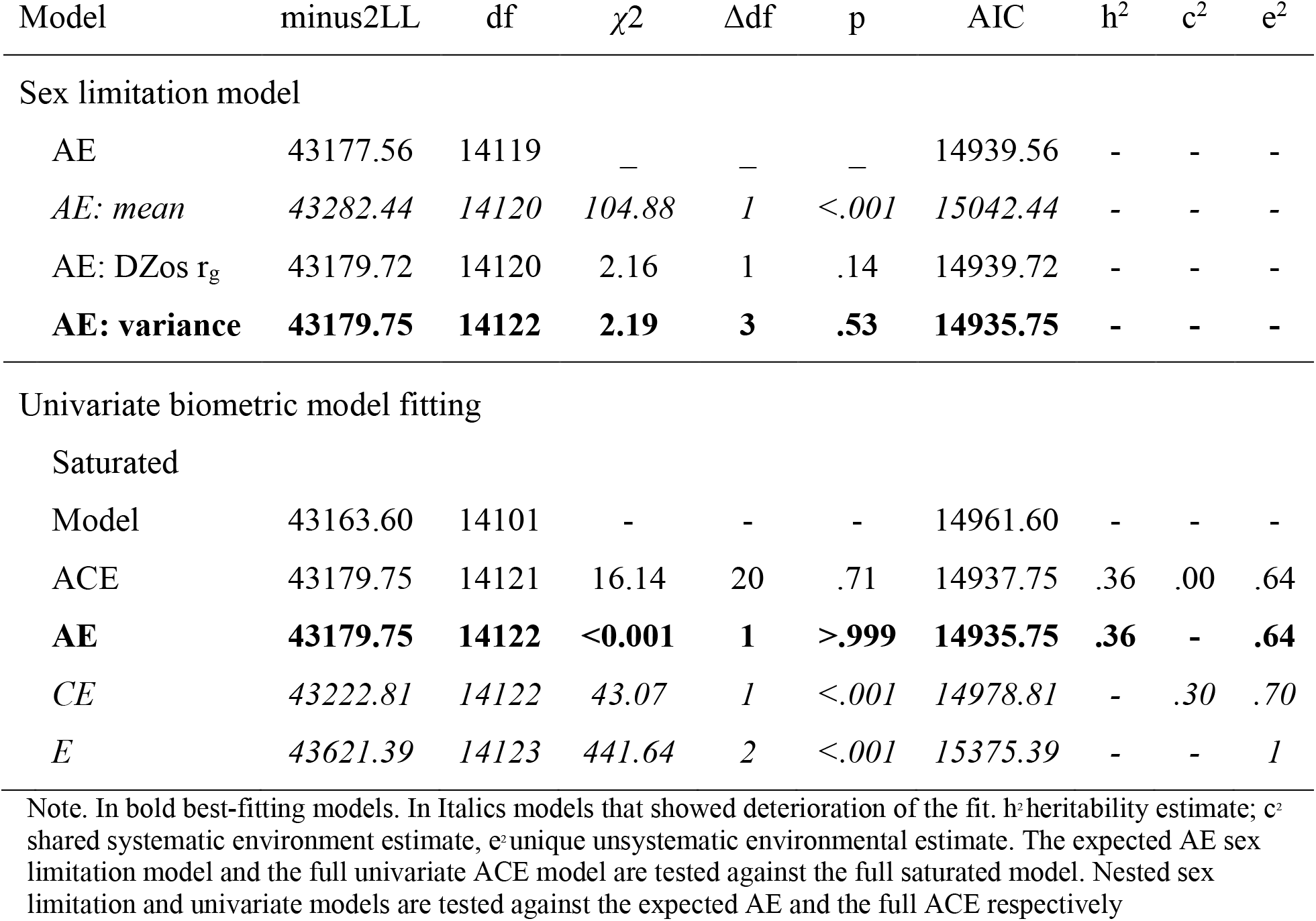
Biometric model: model-fitting results for the sex limitation and the univariate models

| Model | minus2LL | df | $\chi^2$ | $\Delta df$ | p | AIC | $h^2$ | $c^2$ | $e^2$ |
| --- | --- | --- | --- | --- | --- | --- | --- | --- | --- |
| Sex limitation model |  |  |  |  |  |  |  |  |  |
| AE | 43177.56 | 14119 | — | — | — | 14939.56 | - | - | - |
| *AE: mean* | *43282.44* | *14120* | *104.88* | *1* | *<.001* | *15042.44* | - | - | - |
| AE: DZos $r_g$ | 43179.72 | 14120 | 2.16 | 1 | .14 | 14939.72 | - | - | - |
| **AE: variance** | **43179.75** | **14122** | **2.19** | **3** | **.53** | **14935.75** | - | - | - |
| Univariate biometric model fitting |  |  |  |  |  |  |  |  |  |
| Saturated |  |  |  |  |  |  |  |  |  |
| Model | 43163.60 | 14101 | - | - | - | 14961.60 | - | - | - |
| ACE | 43179.75 | 14121 | 16.14 | 20 | .71 | 14937.75 | .36 | .00 | .64 |
| **AE** | **43179.75** | **14122** | **<0.001** | **1** | **>.999** | **14935.75** | **.36** | - | **.64** |
| *CE* | *43222.81* | *14122* | *43.07* | *1* | *<.001* | *14978.81* | - | .30 | .70 |
| *E* | *43621.39* | *14123* | *441.64* | *2* | *<.001* | *15375.39* | - | - | *1* |

Note. In bold best-fitting models. In Italics models that showed deterioration of the fit. $h^2$ heritability estimate; $c^2$ shared systematic environment estimate, $e^2$ unique unsystematic environmental estimate. The expected AE sex limitation model and the full univariate ACE model are tested against the full saturated model. Nested sex limitation and univariate models are tested against the expected AE and the full ACE respectively

Phenotypic correlations, obtained from the most parsimonious model table 2, were *r* = .37 ([.33,.40] 95% CI) within MZ and *r* = .15 ([.11,.19] 95% CI) within DZ twin pairs. Phenotypic correlations suggested once more the AE model as the most appropriate model to describe the data.

The final genetic univariate model fitting results and comparison are presented in table 3. Constraining variance components A, and A and C to zero respectively deteriorated the model fit (all *p =* <.*001*). As expected, the final model AE (Figure 3), with mean estimates adjusted for age (*β*_*age*_ = 0.01) equal to 2.03 for males and 2.25 for females (SD = 1.13), was the most parsimonious well-fitting one (−2LL = *χ*2(1) = **<**0.001, *p = >*.*999*). As shown in table 3, the heritability estimate for the propensity of feeling chills is 36 % (A = .36 [.33, .39] 95% CI), while the remaining 64% of the phenotypic variance (E = .64 [.61, .67] 95% CI) can be accounted for by unsystematic effects, such as environmental experience unique to the individual and measurement errors.

**Figure 3.**
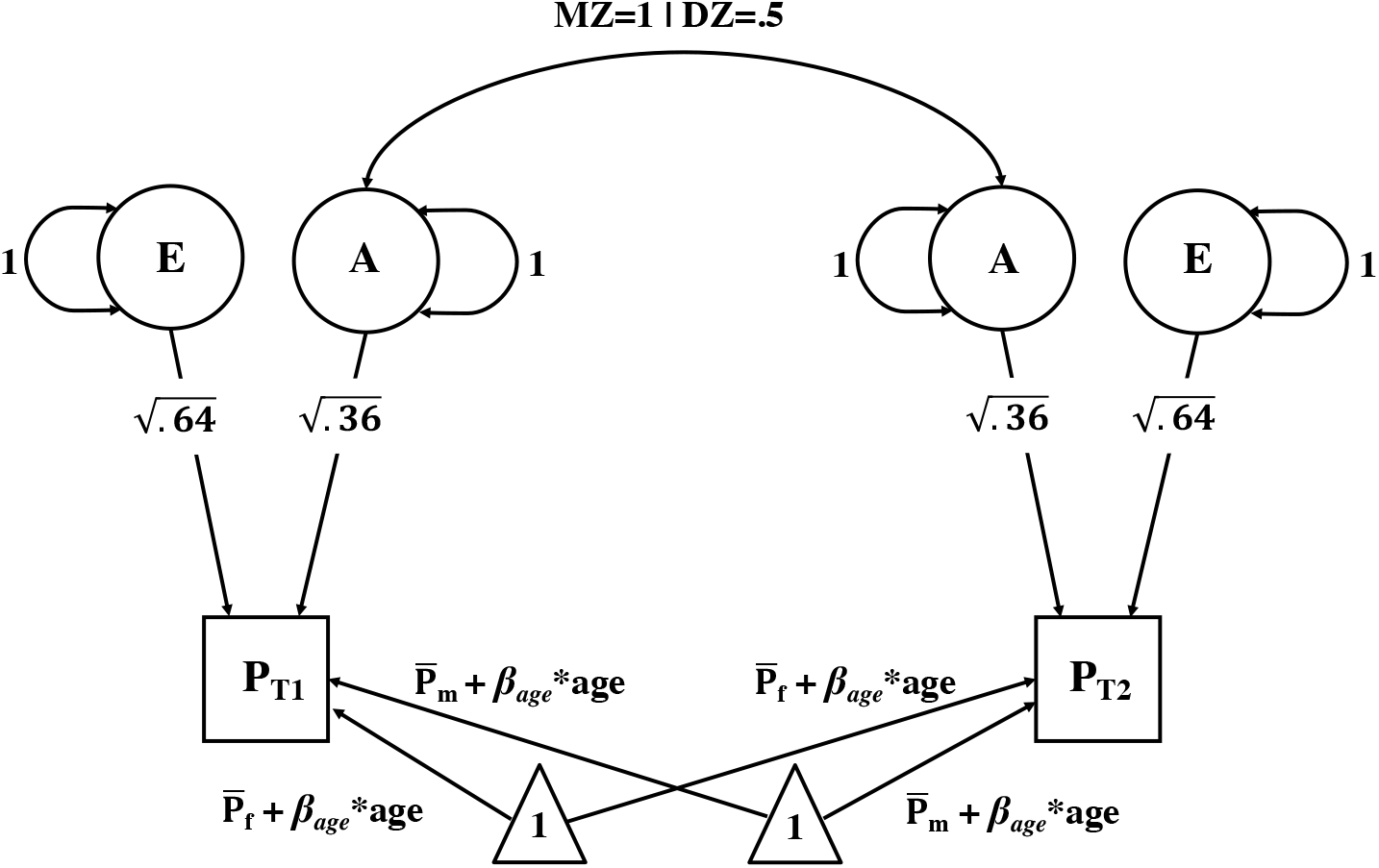
Best fitting AE biometric model. Final model with the five estimated parameters: 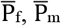, β_0,_ and variance components A and E. The Squares represent twin (T) 1 and 2 observed phenotypic variance (P) in aesthetic chills (with P_T1_= P_T2,_ see table 2) The triangle represents the mean estimates. The circles represent the additive genetic (A) and the environmental (E) variance components, with their associated variance. The arrows pointing to the square represent the genetic and environmental path coefficients. The double arrows across the variance component A represent the expected correlation within MZ (1) and within DZ (.5) twin pairs. 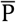 = phenotypic means for females 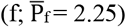 or males 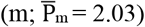; *β*_*age*_ = 0.01. NB for simplicity, we report variance component A and E to be equal to 1, although in our model they were directly estimated (i.e., path coefficients were constrained to be equal to 1)

## Discussion

This research investigates and report genetic and environmental sources of variation for the propensity of feeling aesthetic chills. We analyzed the variance of the NEO-FFI Item 43: “Sometimes when I am reading poetry or looking at a work of art, I feel a chill or wave of excitement”, which is a proxy measurement for the propensity of feeling aesthetic chills. We found that 36% of the variance in feeling aesthetic chills can be explained by additive genetic factors and the remaining 64% by environmental sources of variation. If we consider these results in the view of the test-retest reliability, which we obtained for data from surveys 4, 5 and 9 years apart, which ranged from around 0.52 to 0.61, the environmental variance reflects to a substantial extent reliable non-genetic causes of individual differences.

A lack of shared environmental sources of variation in the propensity of feeling chills should not come as a surprise, given that variation in most of the psychological human traits investigated so far show no relationship with shared environmental effects ^54^. Moreover, we confirmed, in the largest sample used to investigate the role of demographic factors on chills to date, that females and older individuals are more prone than males and younger individuals to experience aesthetic chills. This is in line with some of the results previously obtained from the general population ^15,26,27^. Additionally, it is also important to note that the directionality of the sex effect as found in this study was also consistent, as far as we know, with all of the studies that reported non-significant trends toward females reporting more frequent chills than males^2,29,31^.

We would like to note that, although apparently consistent with the separation call phenomena proposed by Panksepp ^15^, our result are only partially supporting his hypothesis. Panksepp argued that chills are a genetically influenced trait that “resonate with ancient emotional circuits that establish internal social values” ^15^, and that they probably evolved over time from a need for physical closeness induced by the separation between a mother and her infant. Such mother-infant relationships should have produced an enhanced selective propensity of experiencing chills in females. Our finding that variation in chills is influenced by genetic factors, and that females are more prone to experience chills, therefore partially support this hypothesis. However, our findings that no qualitative, nor quantitative, genotype x sex interaction effects affect variation in propensity on aesthetic chills do not support the separation-call hypothesis. Indeed, in line with results on a majority of other human traits ^50,55^, our results are consistent with the hypothesis that the same genetic sources of variation in males and females influence variation in the propensity of feeling aesthetic chills.

We believe our results brings into focus questions that go beyond the descriptive nature of this study. Williams et al. ^34^ argued that the enhanced connectivity between sensory and salience/default networks as found in their study, based on resting-state fMRI data of 1000 subjects from the Human Connectome Project (HCP), indicate that individuals who are more prone to experience chills from art and poetry, as measured by the item 43, are also individuals that can better integrate sensory information with internal emotional experiences. Moreover, preliminary evidence on a small sample recently indicated that augmenting sensory signals that mimics the physical experience of aesthetic chills can enhance individual social-affective cognition (e.g., empathy and pleasure; ^56^).

Yet, how can one reinterpret such results in light of our present findings? Is it environmental exposure over one’s own lifetime to art and poetry that is causally shaping the connectivity as seen in Williams et al. ^34^ or is it more a priori predisposition that makes individuals better at integrating sensory information with their internal states, or alternatively more sensitive to sensory signals such as the ones seen in Haar et al. ^56^, that makes them more likely to reach such peaks of emotional-hedonic experiences? Clearly, further studies are needed to answer these questions. However, our finding that approximately one-third of the variance in the propensity of feeling chills can be explained by genetic influences sheds some light on such questions.

Finally, it is important to consider the limitations of our study. Given that the scope of this research was to explore the etiology of aesthetic chills using an already existing sample of phenotyped individuals, we were constrained by what measurements were available. This limited our capacity to obtain a more nuanced estimate of aesthetic chills. The NEO-FFI item 43 explicitly asks for the propensity of feeling chills only from art or poetry. For example, although a previous study found the item 43 to be correlated with the number of chills experienced from music ^24^, further studies are still needed to claim that etiological sources of variation influence the propensity of feeling chills from music specifically. Moreover, even though the test-retest reliability for the item 43 may be considered good for a single item, it was not perfect. However, if this had any impact on our results it was through increasing the measurement error, which by definition is included in the estimate for the unsystematic environmental sources of variation. As such, our estimates on the effect of additive genetic variation should be considered to be, at worst, a lower bound for the real effect of genetic influence on the propensity of aesthetic chills. Finally, it is important to note that it would be premature to reach any conclusions regarding the putative mechanisms underlying the genetic factors influencing aesthetic chills on the basis of the findings of this study alone. We thus call for further genetic informative studies, such as genome wide association and functional annotation studies, to explore the whole genome in order to inspect which variation, likely polygenic, can be associated with variation in the propensity of aesthetic chills, and to inspect whether variations associated with the propensity of feeling chills are enriched in brain tissue or elsewhere in the body.

## Conclusion

Aesthetic chills are a somatic marker of peak emotional hedonic responses. Previous inconsistent evidence suggested that demographic, and somewhat more consistent evidence has suggested that personality and neurobiological factors, can account for part of the observed variation in feeling chills. Here, we confirmed that females are more prone than males, even if to a small degree, to experience aesthetic chills and that older individuals tend to report experiencing chills more than younger ones. Critically, we revealed that genetics play a role in the individual differences in the propensity of feeling chills, thus indicating that the tendency of experiencing hedonic peaks of emotional reactions to art and poetry is not shaped by nurture alone, but is also influenced by genetic predispositions, as thousands of other human traits are ^54^.

## Acknowledgements

The data reported in this study were collected by the Netherlands Twin Register (NTR, see: www.tweelingenregister.org), a longitudinal cohort established in 1987 by the Department of Biological Psychology at the Vrije Universiteit Amsterdam, registered with the Dutch Data Protection Authority (nr m1412317). The procedure for the formal request of phenotypic data was carried out after the approval of the data access committee, and following the guidelines provided by the NTR. The collection of the data in this study occurred from the year 2004 till the year 2013 ^42^. We thus thank the NTR for their collaboration, and all of the twins that took part in the study. We also thank MacKenzie D Trupp for critical feedbacks on the manuscript, Surabhi S Nath for feedbacks on sections of the code used to analyze the data, and Aenne A Brielmann for advices on how to make this research more open access. This study was partially supported by the BMBF and Max Planck Society. We acknowledge funding for data collection by the Borderline Foundation (USA); ZonMW -31160008 (Netherlands Medical Research), ERC – 284167 (European Research Council) and NWO-480-15-001/674 (Netherlands Organization for Science Research).

## Data Availability Statement

The data that support the findings of this study are available from Netherlands Twin Register (NTR; www.tweelingenregister.org) but restrictions apply to the availability of these data, which were used under license for the current study, and so are not publicly available. Readers interested in having access to the data can consult Ligthart et al. ^42^. The code used to analyze the data is available at https://github.com/giacomobignardi/h2-aesthetic-chills.

## Author contribution

G.B, R.C., and D.I.B. conceived the study; G.B., and S.T.K., analyzed the data; S.T.K. validated the work; Z.T. conceptually validated the work; G.B. visualized the data; G.B. drafted the manuscript; G.B., R.C., Z.T., and D.I.B. edited the manuscript; R.C. and D.I.B supervised the research; D.I.B. was responsible for collecting funding for previous data acquisition relevant for this research. All authors reviewed the last version of this manuscript.

## References

1. McCrae, R. R. Aesthetic Chills as a Universal Marker of Openness to Experience. Motiv. Emot. 31, 5–11 (2007).

2. Grewe, O., Kopiez, R. & Altenmüüller, E. The Chill Parameter: Goose Bumps and Shivers as Promising Measures in Emotion Research. Music Percept. Interdiscip. J. 27, 61–74 (2009).

3. Wassiliwizky, E., Koelsch, S., Wagner, V., Jacobsen, T. & Menninghaus, W. The emotional power of poetry: neural circuitry, psychophysiology and compositional principles. Soc. Cogn. Affect. Neurosci. 12, 1229–1240 (2017).

4. Sumpf, M., Jentschke, S. & Koelsch, S. Effects of Aesthetic Chills on a Cardiac Signature of Emotionality. PLoS ONE 10, (2015).

5. Goldstein, A. Thrills in response to music and other stimuli. Physiol. Psychol. 8, 126–129 (1980).

6. Schurtz, D. R. et al. Exploring the social aspects of goose bumps and their role in awe and envy. Motiv. Emot. 36, 205–217 (2012).

7. Blood, A. J. & Zatorre, R. J. Intensely pleasurable responses to music correlate with activity in brain regions implicated in reward and emotion. Proc. Natl. Acad. Sci. U. S. A. 98, 11818–11823 (2001).

8. Mas-Herrero, E., Zatorre, R. J., Rodriguez-Fornells, A. & Marco-Pallarés, J. Dissociation between Musical and Monetary Reward Responses in Specific Musical Anhedonia. Curr. Biol. 24, 699–704 (2014).

9. Salimpoor, V. N., Benovoy, M., Larcher, K., Dagher, A. & Zatorre, R. J. Anatomically distinct dopamine release during anticipation and experience of peak emotion to music. Nat. Neurosci. 14, 257–262 (2011).

10. Bannister, S. A survey into the experience of musically induced chills: Emotions, situations and music. Psychol. Music 0305735618798024 (2018) doi:10.1177/0305735618798024.

11. Grewe, O., Kopiez, R. & Altenmüller, E. Chills As an Indicator of Individual Emotional Peaks. Ann. N. Y. Acad. Sci. 1169, 351–354 (2009).

12. Mori, K. & Iwanaga, M. Two types of peak emotional responses to music: The psychophysiology of chills and tears. Sci. Rep. 7, 46063 (2017).

13. Rickard, N. S. Intense emotional responses to music: a test of the physiological arousal hypothesis. Psychol. Music 32, 371–388 (2004).

14. Wassiliwizky, E., Wagner, V., Jacobsen, T. & Menninghaus, W. Art-elicited chills indicate states of being moved. Psychol. Aesthet. Creat. Arts 9, 405–416 (2015).

15. Panksepp, J. The Emotional Sources of ‘Chills’ Induced by Music. Music Percept. Interdiscip. J. 13, 171–207 (1995).

16. Sloboda, J. A. Music Structure and Emotional Response: Some Empirical Findings. Psychol. Music 19, 110–120 (1991).

17. Craig, D. G. An Exploratory Study of Physiological Changes during “Chills” Induced by Music. Music. Sci. 9, 273–287 (2005).

18. Grewe, O., Katzur, B., Kopiez, R. & Altenmüller, E. Chills in different sensory domains: Frisson elicited by acoustical, visual, tactile and gustatory stimuli. Psychol. Music 39, 220–239 (2011).

19. Sachs, M. E., Ellis, R. J., Schlaug, G. & Loui, P. Brain connectivity reflects human aesthetic responses to music. Soc. Cogn. Affect. Neurosci. 11, 884–891 (2016).

20. Starcke, K., von Georgi, R., Tiihonen, T. M., Laczika, K.-F. & Reuter, C. Don’t drink and chill: Effects of alcohol on subjective and physiological reactions during music listening and their relationships with personality and listening habits. Int. J. Psychophysiol. 142, 25–32 (2019).

21. Wassiliwizky, E., Jacobsen, T., Heinrich, J., Schneiderbauer, M. & Menninghaus, W. Tears Falling on Goosebumps: Co-occurrence of Emotional Lacrimation and Emotional Piloerection Indicates a Psychophysiological Climax in Emotional Arousal. Front. Psychol. 8, (2017).

22. Herman, A. M., Palmer, C., Azevedo, R. T. & Tsakiris, M. Neural divergence and convergence for attention to and detection of interoceptive and somatosensory stimuli. Cortex 135, 186–206 (2021).

23. Klepzig, K. et al. Brain imaging of chill reactions to pleasant and unpleasant sounds. Behav. Brain Res. 380, 112417 (2020).

24. Colver, M. C. & El-Alayli, A. Getting aesthetic chills from music: The connection between openness to experience and frisson. Psychol. Music 44, 413–427 (2016).

25. Nusbaum, E. C. & Silvia, P. J. Shivers and Timbres: Personality and the Experience of Chills From Music. Soc. Psychol. Personal. Sci. 2, 199–204 (2011).

26. Balte?, F.R. & Miu, A. C. Emotions during live music performance: Links with individual differences in empathy, visual imagery, and mood. Psychomusicology Music Mind Brain 24, 58–65 (2014).

27. Benedek, M. & Kaernbach, C. Physiological correlates and emotional specificity of human piloerection. Biol. Psychol. 86, 320–329 (2011).

28. Mori, K. & Iwanaga, M. Resting physiological arousal is associated with the experience of music-induced chills. Int. J. Psychophysiol. 93, 220–226 (2014).

29. Bannister, S. Distinct varieties of aesthetic chills in response to multimedia. PLOS ONE 14, e0224974 (2019).

30. Guhn, M., Hamm, A. & Zentner, M. Physiological and Musico-Acoustic Correlates of the Chill Response. Music Percept. Interdiscip. J. 24, 473–484 (2007).

31. Silvia, P. J. & Nusbaum, E. C. On personality and piloerection: Individual differences in aesthetic chills and other unusual aesthetic experiences. Psychol. Aesthet. Creat. Arts 5, 208–214 (2011).

32. Silvia, P. J., Fayn, K., Nusbaum, E. C. & Beaty, R. E. Openness to experience and awe in response to nature and music: Personality and profound aesthetic experiences. Psychol. Aesthet. Creat. Arts 9, 376–384 (2015).

33. Mori, K. & Iwanaga, M. General reward sensitivity predicts intensity of music-evoked chills. Music Percept. 32, 484–492 (2015).

34. Williams, P. G., Johnson, K. T., Curtis, B. J., King, J. B. & Anderson, J. S. Individual differences in aesthetic engagement are reflected in resting-state fMRI connectivity: Implications for stress resilience. NeuroImage 179, 156–165 (2018).

35. Barron, F. & Parisi, P. Twin Resemblances in Creativity and in Esthetic and Emotional Expression. Acta Genet. Medicae Gemellol. Twin Res. 25, 213–217 (1976).

36. Barron, F. Twin resemblances in creative thiking and aesthetic judgment. In Artists in the making. (1972) New York, NY: Seminar Press., (pp. 174–181).

37. Butkovic, A., Ullén, F. & Mosing, M. A. Personality related traits as predictors of music practice: Underlying environmental and genetic influences. Personal. Individ. Differ. 74, 133–138 (2015).

38. Germine, L. et al. Individual Aesthetic Preferences for Faces Are Shaped Mostly by Environments, Not Genes. Curr. Biol. CB 25, 2684–2689 (2015).

39. Sutherland, C. A. M. et al. Individual differences in trust evaluations are shaped mostly by environments, not genes. Proc. Natl. Acad. Sci. 117, 10218–10224 (2020).

40. Zietsch, B. P., Lee, A. J., Sherlock, J. M. & Jern, P. Variation in Women’s Preferences Regarding Male Facial Masculinity Is Better Explained by Genetic Differences Than by Previously Identified Context-Dependent Effects. Psychol. Sci. 26, 1440–1448 (2015).

41. Costa, P. T., Jr. & McCrae, R. R. The Revised NEO Personality Inventory (NEO-PI-R). in The SAGE Handbook of Personality Theory and Assessment: Volume 2 — Personality Measurement and Testing 179–198 (SAGE Publications Ltd, 2008). doi:10.4135/9781849200479.

42. Ligthart, L. et al. The Netherlands Twin Register: Longitudinal Research Based on Twin and Twin-Family Designs. Twin Res. Hum. Genet. 1–14 (2019) doi:10.1017/thg.2019.93.

43. Distel, M. A. et al. Personality, health and lifestyle in a questionnaire family study: a comparison between highly cooperative and less cooperative families. Twin Res. Hum. Genet. Off. J. Int. Soc. Twin Stud. 10, 348–353 (2007).

44. Geels, L. M. et al. Increases in alcohol consumption in women and elderly groups: evidence from an epidemiological study. BMC Public Health 13, 1–13 (2013).

45. Treur, J. L., Boomsma, D. I., Ligthart, L., Willemsen, G. & Vink, J. M. Heritability of high sugar consumption through drinks and the genetic correlation with substance use. Am. J. Clin. Nutr. 104, 1144–1150 (2016).

46. Boomsma, D. I., Busjahn, A. & Peltonen, L. Classical twin studies and beyond. Nat. Rev. Genet. 3, 872–882 (2002).

47. Ouwens, K. G. et al. A characterization of postzygotic mutations identified in monozygotic twins. Hum. Mutat. 39, 1393–1401 (2018).

48. Jonsson, H. et al. Differences between germline genomes of monozygotic twins. Nat. Genet. 53, 27–34 (2021).

49. Knopik, V. S., Neiderhiser, J. M., DeFries, J. C. & Plomin, R. Behavioral Genetics. (Worth Publishers, 2016).

50. Vink, J. M. et al. Sex Differences in Genetic Architecture of Complex Phenotypes? PLOS ONE 7, e47371 (2012).

51. Boker, S. M. et al. OpenMx: Extended Structural Equation Modelling. (2019).

52. Pritikin, J. N., Hunter, M. D. & Boker, S. M. Modular Open-Source Software for Item Factor Analysis. Educ. Psychol. Meas. 75, 458–474 (2015).

53. Hunter, M. D. State Space Modeling in an Open Source, Modular, Structural Equation Modeling Environment. Struct. Equ. Model. Multidiscip. J. 25, 307–324 (2018).

54. Polderman, T. J. C. et al. Meta-analysis of the heritability of human traits based on fifty years of twin studies. Nat. Genet. 47, 702–709 (2015).

55. Stringer, S., Polderman, T. J. C. & Posthuma, D. Majority of human traits do not show evidence for sex-specific genetic and environmental effects. Sci. Rep. 7, 8688 (2017).

56. Haar, A. J. H., Jain, A., Schoeller, F. & Maes, P. Augmenting aesthetic chills using a wearable prosthesis improves their downstream effects on reward and social cognition. Sci. Rep. 10, 21603 (2020).

